# A miniature kinematic coupling device for mouse head fixation

**DOI:** 10.1101/2021.10.26.463065

**Authors:** Su Jin Kim, Alexander H. Slocum, Benjamin B. Scott

## Abstract

Head-fixation is a common technique in the preparation of subjects for neuroscience experiments. Accurate alignment, stability, and repeatability during fixation provide experimental consistency, thus enabling the subject to return to the same position over time to provide meaningful data. Head restraint systems inspired by kinematic clamps have been developed to allow micron scale repositioning across imaging epochs in rats. Here we report the development of a light-weight, implantable kinematic coupling (clamp) system that is wearable by mice, and enables repeated positioning to submicron accuracy across imaging epochs. This system uses a stainless steel headplate and a Maxwell-style three-groove kinematic mounting system with magnetic force clamping load. Spheres on the dorsal surface of the headplate provide contact points for vee-groove kinematic features machined into a tabletop mount. Evaluation of the clamp using multiphoton microscopy revealed submicron precision in registration accuracy and stability, allowing cellular resolution calcium imaging in awake, behaving mice. These results indicate that miniaturized implantable kinematic clamps for mice could be valuable for future experiments which require repositioning of subjects across time and different instruments.

**Highlights:**

- Development of a kinematic clamp for mice for precise repositioning in chronic studies.
- Headplate and clamp provide stability for cellular resolution imaging during behavior.
- Ruby contact features enable submicron registration repeatability.

## Introduction

Head-fixation is widely used in neuroscience for stimulus control, behavioral monitoring, and neural recording and perturbation experiments, because it minimizes motion between the subject’s head and the recording or stimulating equipment. Experiments which involve repeated use of the same subject, such as developmental or chronic studies, use mechanical restraint systems to facilitate alignment and stability between the head and experiment apparatus (Noda *et al*., 1971, Populin and Yin 1998, Tratchenberg *et al*., 2002, Holtmaat 2009). Accurate and reproducible alignment is valuable in experimental pipelines in which the same subject is transferred across instruments, thus requiring adherence to a common coordinate system (de Vries *et al*., 2020). Cellular resolution optical imaging (Denk *et al*., 1990) is among the *in vivo* recording technologies that require high stability and repositioning accuracy, typically on the micron scale (Dombeck *et al*., 2007).

Several head fixation systems have been designed which achieve the mechanical stability necessary for imaging and enable reliable repositioning within tens of microns. To achieve even higher registration accuracy, Tank and colleagues developed a head-fixation system for rats inspired by kinematic clamps (Scott *et al*., 2013 and Rich *et al*., 2018). Kinematic clamps (Maxwell 1890, Evans 1989) are widely used in optical and mechanical devices that require precise alignment. These coupling systems achieve precise (repeatable) repositioning by constraining each of the three axes of translation (X, Y, Z) and rotation (yaw, pitch, roll) of object movement, and by applying the minimum number of constraints (Smith and Chetwynd 1992, Slocum 1992a, for a review see Slocum 2010). Such head fixation systems allow for micron-scale stability and registration accuracy. By combining implantable kinematic clamps with operant training systems, Tank, Brody and colleagues developed an automated imaging system in which rats were trained to submit to brief periods of restraint while performing a perceptual decision-making task (Scott *et al*., 2015, 2017). The high accuracy and stability of the registration system allowed the same neurons to be recorded on each imaging epoch.

Voluntary head restraint systems for rodents were first used to facilitate presentation of sensory stimuli to the same region of visual space (Girman 1980, Girman 1985). Reports that rats could be trained to self head-fix using computer-controlled training systems (Kampff *et al*., 2010) inspired a resurgence of technological advancements for voluntary head restraint. Since then, several systems that involve voluntary head fixation in mice have been developed (Murphy *et al*., 2016, Aoki *et al*., 2017, Murphy *et al*. 2020, Hao *et al*., 2021), however to our knowledge none have demonstrated micron-scale repositioning accuracy necessary for cellular resolution imaging.

Here we describe the development and evaluation of a kinematic clamp head restraint system for mice. Our goal was to achieve a headplate light enough to be worn by a mouse, but stable enough for multiphoton imaging during behavior, with repositioning accuracy on the micron-scale or better. Scott *et al*., used a Kelvin-style kinematic system based on a groove, socket (which should be a trihedron), and flat to engage spherical surfaces. The three-groove system favored by Maxwell is simpler to manufacture—which potentially enables greater performance (Slocum 1992c)—and provides an effective center of rotation at the center of the coupling, which in this case would be the neurological point of interest. We evaluated the performance of several variants of the clamp fabricated from different contact materials. We show that kinematic head fixation systems based on the three-groove (i.e., Maxwell) design with ruby contact features achieved position repeatability in the submicron range. Proof-of-principle imaging experiments demonstrate that such a clamp design could be integrated into an imaging system for voluntary restraint system or for time lapse studies, in which the same neuronal populations are tracked over development or learning.

## Methods

### Fabrication of the coupling system

Headplates and mounts were designed using CAD software (SolidWorks, Dassault Systemes) guided by the publicly available kinematic coupling design spreadsheet (Slocum 2016). The mount was made from milled 7075 aluminum (RA 53.5, Protolabs). Neodymium magnets were seated on the top (dorsal) surface in square-shaped depressions, symmetric in position to each vee-groove. Axially magnetized magnets (B662-N52, 6.4lb magnetic attraction force, K&J Magnetics) were bonded using structural epoxy adhesive (Loctite E-05MR, Henkel).

Headplates were fabricated from 17-4 stainless steel (RC 33, Protolabs). 17-4 stainless steel is not as hard as 440 stainless steel, but has superior corrosion resistance, and is magnetic, hence it was chosen. Ruby spheres (0.1250” diameter, G25, Mohs Hardness 9, Bird Precision), and 304 stainless steel bearings (0.125” diameter, G100, <28 Hardness Rockwell C, uxcell, Amazon) were each tested and located and attached in conical depressions on the top (dorsal) surface of the headplate with epoxy (Loctite E-05MR, Henkel) and left to cure overnight.

### Assessment of surface finish and smoothness with Scanning electron microscopy

Scanning electron microscopy was conducted with a Supra 55VP instrument operated at 3 kV. The angle of the scanning beam with respect to the sample surface was 90°. Prior to imaging, the headplates and the clamp were cleaned with acetone (Sigma-Aldrich). Headplate samples were affixed to a stub and photographed at 30× and 2,000× magnification. Vee groove samples were photographed at 50× and 2,000× magnification. Each sample was photographed under similar conditions and working distance (10-11 mm for headplate samples, 7-8 mm for the vee groove sample) to allow for comparisons. SEM photos were digitized using SmartSEM (Zeiss).

### Assessment of surface finish and smoothness with Optical profiler

The clamp feature surface finishes were assessed using a noncontact white light interferometer (ZYGO NewView 9000). The headplates were placed under the optical profiler objective for 3-dimensional scanning. To separate roughness from the shape of the surface, we subtracted a spherical shape from the ruby and machined spheres raw data with Mx™ software (ZYGO).

### Assessment of holding force with pull-spring scale

Clamp holding force for the three different ball-groove interfaces was measured using a pull-spring scale (80000042, OHAUS). The clamp was mounted on the edge of an optical breadboard (Thorlabs) using threaded clamping posts (1.5” diameter, P4, Thorlabs). A custom 3D printed adapter with a thru hole (0.5” diameter) was attached to the bottom of each headplate with glue (Loctite 401) and left to dry. The headplate was inserted into the clamp, and the S-hook at the end of the pull-spring scale was hooked in the thru hole attached to the headplate. For each measurement, the pull-spring scale was pulled directly down until the headplate was removed from the clamp, and the force was recorded in Newtons (N). This was repeated five times, and we report the average values: ruby headplate 0.9N, bearing headplate 2.4N, machined headplate 3.4N. The different effective clamp holding forces provided by the magnets are attributed to the different magnetic circuit properties realized by the different ball-groove interfaces.

### Measurement of registration accuracy using two-photon microscopy

Registration accuracy was assessed by optical imaging of fixed fluorescent samples mounted to the kinematic headplates. Samples were composed of a solution of 1μm diameter polystyrene beads (1μm green, Sigma Aldrich) in hard drying fluorescent mounting medium (PolyMount, 18606, PolySciences) at a dilution of 1:80 (beads:mounting medium). Samples were mounted between two glass coverslips (S17521, Fisherbrand), left to dry for 48 hours, then bonded to the headplates with adhesive (Loctite 401, Henkel). Glass coverslips were carefully trimmed to the shape of the headplate using a scoring tool (B01N4AGJ2F, Vivian, Amazon). The kinematic clamp was mounted on an air table (Newport) using threaded clamping posts (1.5” diameter, P4, Thorlabs).

#### Optical measurement of axial position

To measure position along the perpendicular to the imaging plane of the microscope, we bonded a microprism with a silver coated hypotenuse (5.0mm A1, 4531-0029, Tower Optical Corp.) to the top of the kinematic mount, using optical adhesive (Norland NO61). We then bonded a glass coverslip (S17521, Fisherbrand) to the headplates with optical adhesive. Two pieces of glass (Window Rect N-BK7, 4540-0419, Tower Optical Corp) were used to create the sample--one parallel to the headplate to add height, then another perpendicular to the headplate for our sample (see Figure 4A). We trimmed a piece of glass coverslip to the size of the perpendicular piece of glass and fixed a fluorescent bead sample to the perpendicular piece of glass using the same procedure as described in the optical measurement of lateral position. The system was configured such that once the headplate was placed in the mount, the fluorescent sample was reflected on the face of the microprism’s hypotenuse.

The fluorescent beads were imaged using a commercial two-photon (2P) microscope (Bruker Corporation, Prairie View software) with a 20x objective (#46-145, Mitutoyo) at 920nm and laser power 37.5mW.

Registration accuracy was measured by manually inserting the headplate into the clamp. Each insertion consisted of:

1. manually placing the headplate under the clamp such that the kinematic features were roughly aligned with the headplate spheres, approximately 3-5mm away,
2. releasing the headplate and allowing the magnetic force and kinematic features to guide its placement,
3. acquiring a single image to record the position of the fluorescent beads,
4. removing the headplate.

This process was repeated 100 times for lateral position measurements and 30 times for axial position measurements. We define position as the location of the beads relative to the starting location, and displacement as the difference in position between any position and the subsequent position.

#### Assessment of measurement drift

For each imaging session, we assessed baseline variability in the position of the fluorescent sample before and after measuring clamp repositioning accuracy. These measurements were used to define the variability in the position of the headplate that was not due to the kinematic registration system. Such variability could be due to small movements in the position of the microscope objective relative to the table. Keeping the headplate fixed in the clamp, we recorded a Z-stack (0.5μm steps, -10μm to 10μm) and a time series (1 image every minute, for 30 minutes total). This procedure was performed before each registration measurement. Once we identified a period where the variability was < 1.5μm root mean square (RMS) for both directions in the focal plane, we proceeded with the registration measurement session. Bead displacement in the focal plane (X and Y dimensions) was estimated using the sub-pixel rigid-motion correction algorithm NoRMCorre (Pnevmatikakis and Giovannucci 2017). NoRMCorre uses 2D cross-correlation calculations between a reference image and subsequent images taken at each insertion and selects X and Y translations that produce peak correlation value.

#### Assessment of measurement noise

Variability in position estimated by subpixel motion correction may be contaminated several factors including measurement noise (Thompson *et al*., 2002). To assess the noise in our measurements, we attached imaged fluorescent bead samples on the kinematic clamp fixed to optical table. We commanded the microscope to move in steps in the vertical and horizontal directions and took images after each step. Using subpixel motion correction algorithms, we estimated the image displacement and compared this to the known displacement introduced by the microscope motorized stage. We found noise across successive images without stage movement or activating the clamp, to be on the order of 100nm, and found we were able to resolve displacements in position on the order of 0.25μm (Supplemental Figure 1).

### In vivo testing

Adult mice (Thy-1 GCaMP6f, n=2) were habituated to a wheel while head-fixed for one week. Mice were given a 0.2mL sweetened condensed milk reward (milk:water 1:2 dilution) for 30 minutes of walking. Wheel turning was measured using a rotary encoder (H1, US Digital) and an Arduino board (Mega 2560 Rev3, Arduino). Infrared video was used to synchronize imaging data to wheel speed. After training, we assessed brain motion in head-fixed animals using a commercial 2P microscope (Bruker Ultima Investigator) using a 16X water immersion objective (Nikon), and resonant galvo scanning at 30 fps. For calcium imaging we used 920nm and laser power 50mW. For isosbestic point imaging (Barnett *et al*., 2017) we used 820nm and laser power 70mW. We categorized significant calcium events as changes in fluorescence greater than 5 times the standard deviation of the whole trace for 820nm, and changes in fluorescence greater than 5 times the standard deviation of the baseline period (<0.06 ΔF) for 920nm.

### Surgery

All experiments and procedures were performed in accordance with protocols approved by the Boston University Animal Care and Use Committee. Headplate and cannula surgery was performed on healthy male and female GCaMP6f mice. Preoperative injections of dexamethasone (3mg/kg, S.C.) were administered before surgery. Mice were induced with 3% isoflurane, then placed in a stereotaxic frame (KOPF Instruments) and anesthetized with 1.5-2% for surgery. Hair on the scalp was trimmed with scissors, then removed with Nair. A midline incision was made in the scalp, and the skin was retracted to expose the skull. Connective tissue was removed from the dorsal surface of the skull. The dorsal surface of the skull was cleaned with a cotton-tipped applicator soaked in isopropyl alcohol, and scored using a scalpel, to assist with anchoring. The headplate was positioned centered to the cannula and affixed with dental cement (C&B Metabond, Parkell, Edgewood, NY). Optical access was achieved by cortical excavation followed by implantation of a glass bottomed cannula above hippocampus. Animals were assessed one, two, and seven days after surgery for health and window clarity.

## Results

### Design of a miniature kinematic coupling system for mice

We designed our coupling device on the principles of kinematic clamps (Figure 1) where there are six points of contact between two bodies and hence six degrees of freedom can be exactly constrained. We based our designs on Maxwell-style kinematic couplings, in which three spherical surfaces mate with three vee-grooves oriented 120**°** apart, and where the vees are oriented along the angle bisectors of the triangle whose vertices are the sphere’s centers. Maxwell coupling systems (three vees) have improved performance and are known to be more repeatable than the Kelvin-style clamps (vee, flat, cone), which were used in previous head-fixation experiments in rats (Slocum 1995, Evans 1989; Scott et al. 2013). In addition, we designed our system to accommodate subject biology in several ways. First, the headplate needed to be light; 2 grams is the weight well tolerated by mice in previous behavioral studies (Voigts *et al*., 2013). Second, we designed the system to use magnetic clamping force (Rich *et al*. 2018, SfN, abstract). Finally, the headplate needed to be made from biocompatible materials to allow for chronic implantation.

**Figure 1.**
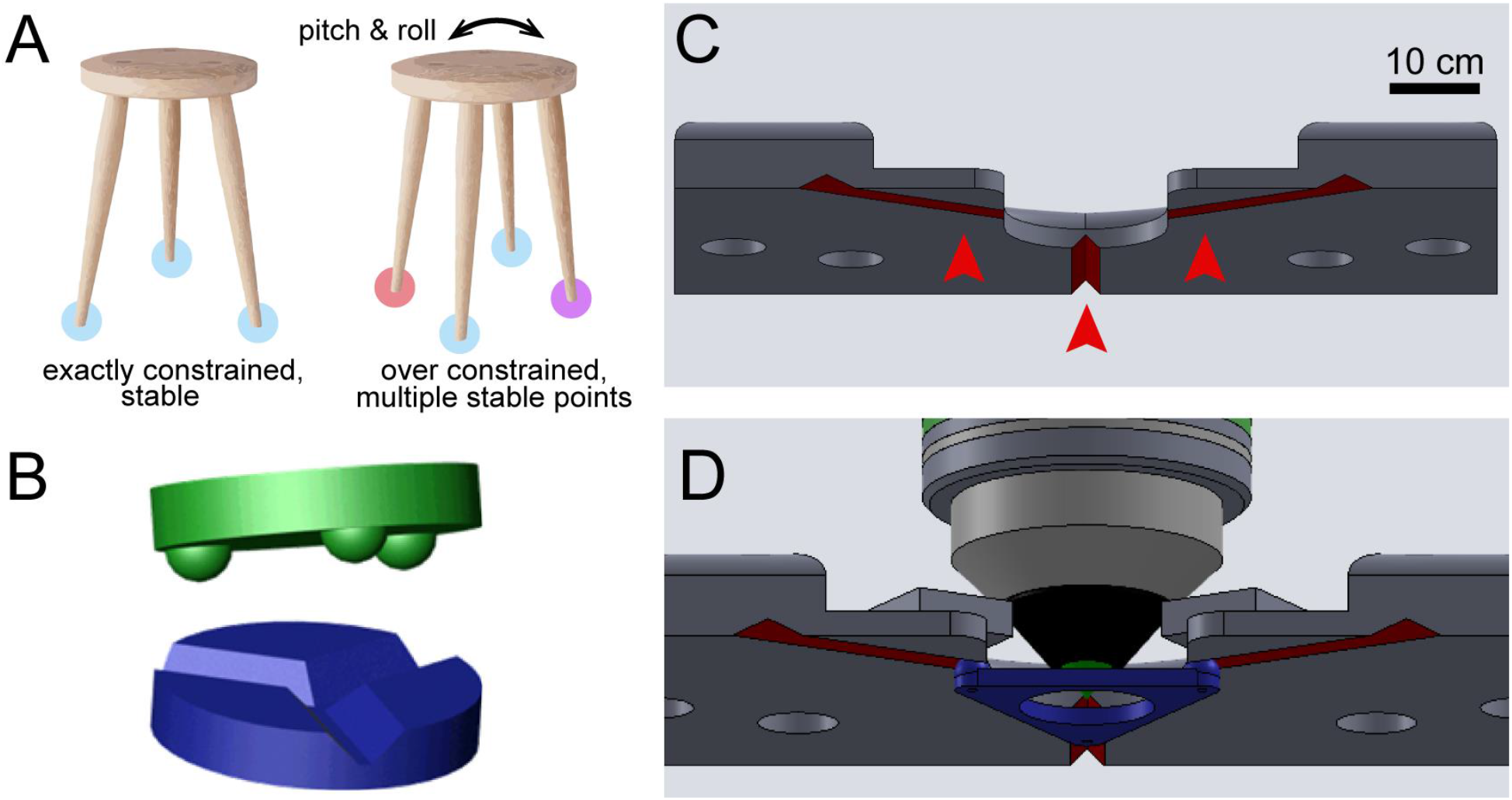
Principle of kinematic coupling and design of miniature clamp for mice. A) Principles of kinematic coupling illustrated by comparing 3- and 4-legged stools. The 3-legged stool (left) has exactly one stable configuration. In contrast, the four-legged stool (right) is susceptible to wobble (pitch and roll), because any three of its legs can create a stable plane of contact with the ground. B) Illustration of a Maxwell clamp, or a three-groove coupling. The three vee grooves, located on the base (blue), are oriented towards the center. Each of the vee grooves provides two contact points, for a total of six contact points, enough to constrain all six degrees of freedom. The mating part (green) has three curved surfaces that seat into the three vee grooves. C) Illustration of a kinematic mount designed to mate with the headplate in B. The mount has a center aperture to accommodate small working distance microscope objectives. Three machined vee grooves (red arrows), oriented 120 degrees apart, provide contact points for the headplate spheres. The inner walls of the vee grooves are angled at 90 degrees. The mount has magnets opposite of the vee grooves to provide clamp force. The mount also has holes for mounting on an optical table. D) Illustration of the kinematic clamp engaged and microscope objective in place.

Headplates were equilateral triangles, 1.5mm thick and 25.4mm long on each side, made from biocompatible magnetic stainless steel (17-4 ph). In the center of the headplate was an 11.9mm diameter aperture to facilitate implantation on the mouse skull and optical access during imaging. We fabricated two variants that differed in the material used for the spherical kinematic features. The first headplate was designed to be machined from a single piece of stainless steel with half-spheres machined into the vertices of the triangular headplate. The second headplate design had conical depressions milled into the headplate corner vertices and stainless steel ball bearings were bonded to the headplate at these depression sites.

For repeatability, the material below the contact point regions must not yield. For metals, this would be shear below the surface at a depth about 1/3^rd^ the diameter of the contact zone diameter. For “infinite” life, the ratio of contact shear stress to material shear yield stress should be less than 0.5 (see Table 2).

### Characterization of clamp contact materials

Given that surface finish is one of the major contributors to repeatability (Slocum and Donmez 1988) we sought to assess the kinematic features by scanning electron microscopy (SEM). SEM imaging revealed the presence of ridges left by the milling process on the machined headplate and mount (Figure 2). While the stainless steel bearings appeared smoother than the machined spheres, they still showed some surface texture which could influence the registration accuracy by providing multiple micro contact points (Figure 2A-F). These data motivated us to construct a third headplate designed with polished ruby spheres, which are ultra-hard and wear resistant, in place of stainless steel bearing balls. Using an optical profiler, we confirmed that the ruby spheres RMS surface roughness was significantly better than the machined spheres, 0.100μm and 0.533μm respectively and in agreement with the bearing grades, which defines their geometric tolerance, provided by the manufacturer. The weights of each headplate are as follows: ruby, 2.2g; bearing, 2.2g; machined half-sphere, 2.1g.

**Figure 2.**
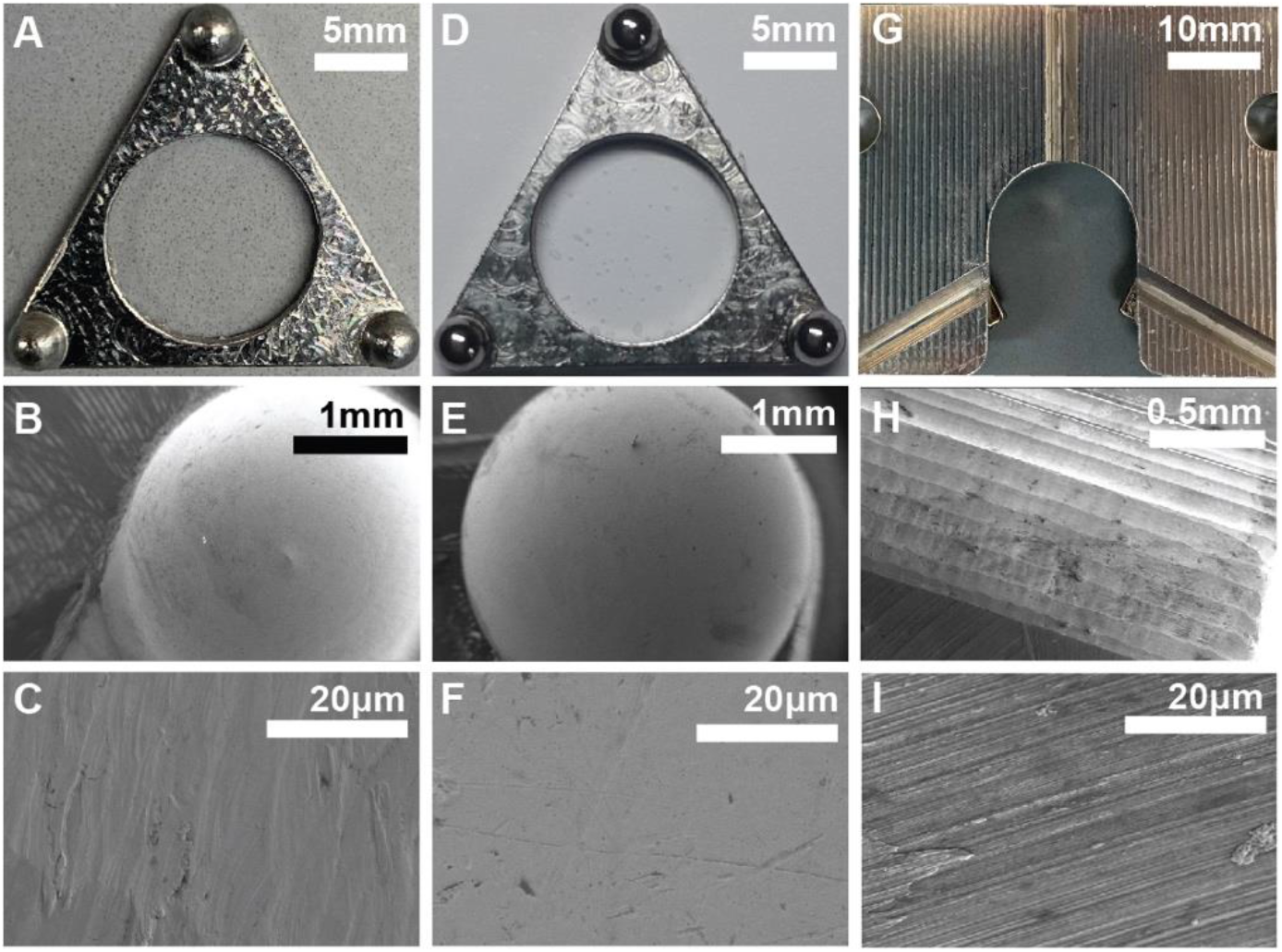
Characterization of kinematic contact features. A) Image of a machined half-sphere headplate. B-C) SEM images of the machined half-sphere. D) Image of a stainless steel bearing ball headplate. E-F) SEM images of the stainless steel bearing ball. G) Image of the machined vee grooves on the Maxwell mount. H-I) SEM images of the machined vee grooves.

### Registration accuracy of clamp contact materials

We next tested the registration accuracy of each of the three headplate designs (machined half sphere, bearing ball, and ruby sphere) using two photon (2P) microscopy (Figure 3). In order to resolve displacement less than the theoretical limit of our 2P microscope, we used sub-pixel motion correction (Pnevmatikakis and Giovannucci, 2017) combined with high signal-to-noise imaging of single beads (Thompson *et al*., 2002). For each headplate we attached a fluorescent bead sample and took images after repeated insertions of the headplate into the clamp (Table 1; Figure 3A-C). The average RMS lateral position for each headplate was as follows: ruby horizontal 0.45μm, ruby vertical 0.65μm, bearing horizontal 3.06μm, bearing vertical 3.74μm, machined horizontal 2.88μm, machined vertical 3.66μm (Figure 3D, for displacement see Figure 3E, 3F). The ruby headplate had the smallest RMS lateral position (Wilcoxon rank sum test, p<0.01 for horizontal and vertical position).

**Figure 3.**
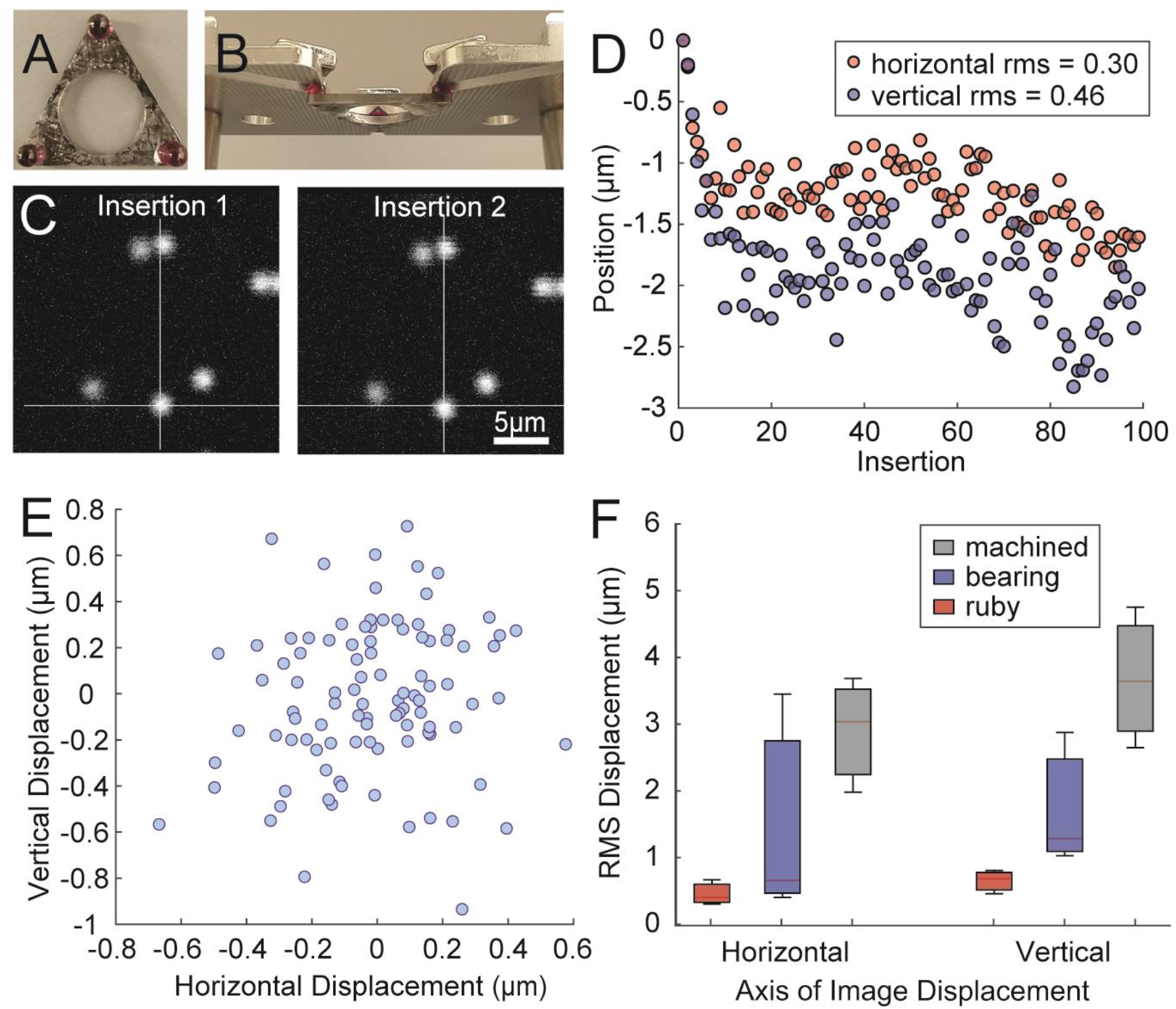
Lateral registration accuracy of the kinematic clamp. A) Image of the headplate with ultrahard ruby kinematic features. B) Image of clamp mated with the headplate. C) Image of 1μm diameter fluorescent spheres mounted to the headplate, obtained with 2P microscopy. Second panel shows the image acquired on a successive activation of the clamp. Comparison of the position of the beads across successive clamp activations was used to estimate registration accuracy. D) Plot showing the position of the beads in the horizontal and vertical axes across successive insertions (horizontal RMS = 0.30μm, vertical RMS = 0.46μm). E) Plot showing location of the beads in X and Y from one session. F) Horizontal and vertical displacement across all headplate contact surfaces.

**Table 1.** Registration accuracy and clamp failure rate of different headplate contact materials.

| **Material** | **rms X**<br>**(microns)** | **rms Y**<br>**(microns)** | **rms Z**<br>**(microns)** | **Clamp**<br>**Failure**<br>**Rate** |
| --- | --- | --- | --- | --- |
| Ruby | 0.3 | 0.46 | 0.02 | 0.02 |
| Ruby | 0.4 | 0.8 | 0.06 | 0.04 |
| Ruby | 0.66 | 0.68 | 0.03 | 0.05 |
| Bearing | 0.4 | 1.02 | 0.16 | 0.01 |
| Bearing | 3.43 | 2.86 | 0.16 | 0 |
| Bearing | 5.37 | 7.34 | 0.59 | 0 |
| Machined | 1.97 | 2.63 | 0.05 | 0.17 |
| Machined | 3.02 | 3.62 | 0.06 | 0.15 |
| Machined | 3.66 | 4.73 | 0.07 | 0.06 |

To assess axial (Z dimension) registration accuracy we used a miniature mirror to bend the microscope’s light path 90 degrees (Figure 4A). We measured X-Y displacements of the imaging plane as a proxy for axial displacements of the headplate following clamp activation (Figure 4B-C). Using this approach, we found a similar pattern observed in the lateral registration accuracy measurements; the ruby headplate had the smallest RMS z-axis displacement (Wilcoxon ranked sum test, p<0.01), followed by the stainless steel bearings, and the machined spheres (0.4μm, 0.8μm, 1.4μm respectively).

**Figure 4.**
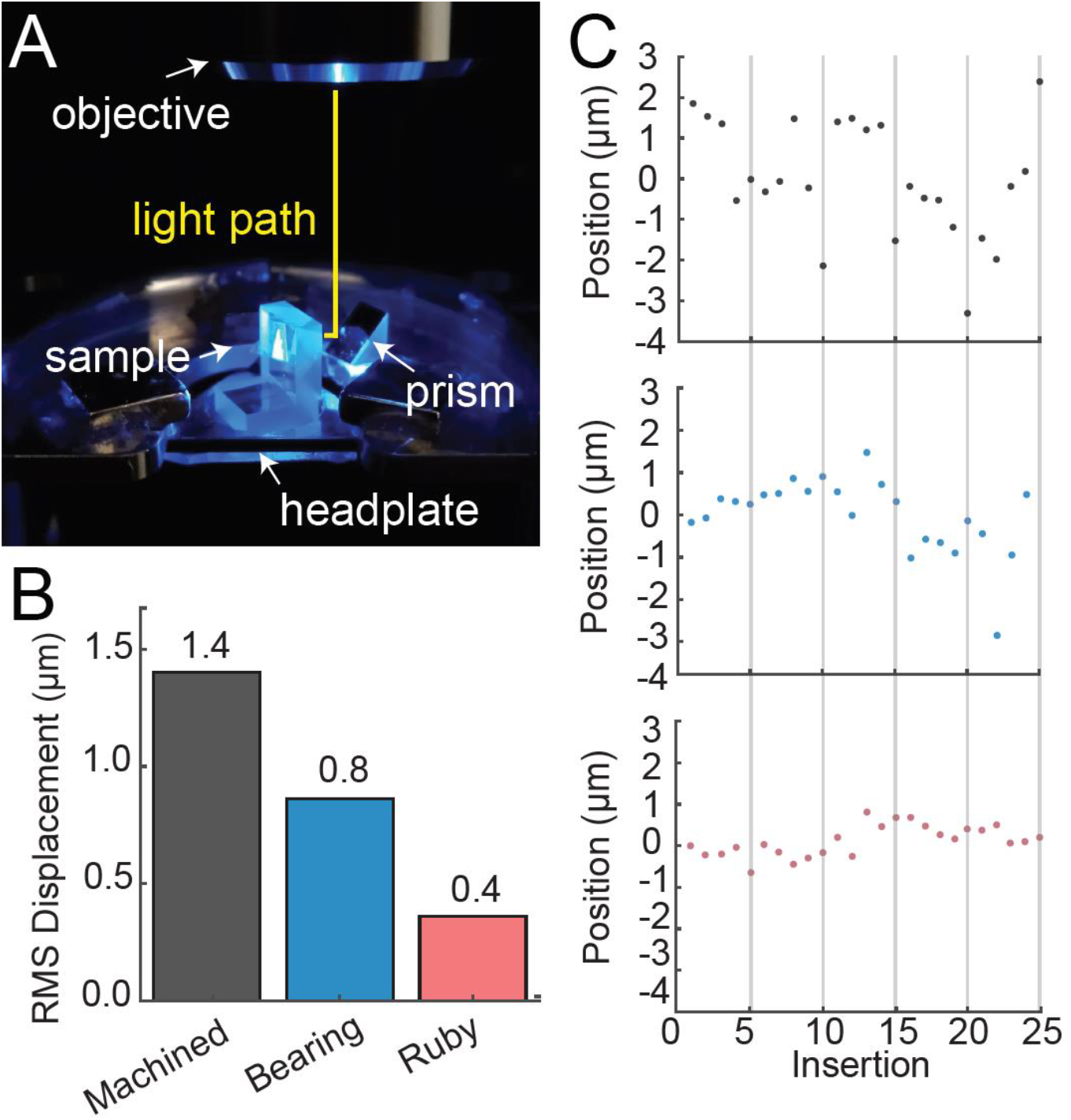
Axial registration accuracy of the kinematic clamp. A) Photograph of the approach used to measure z-axis registration accuracy. A sample comprised of miniature fluorescent spheres was mounted perpendicular to the headplate and a prism with a silvered hypotenuse was used to reflect the light path (yellow line) from the microscope 90 degrees. B) Z-axis position across all headplate contact surfaces (ruby vs. stainless bearing vs. machined half-spheres). C) Plot showing the position of the beads in the Z-axis after each clamp activation for each headplate (top panel, machined; middle panel, bearing; lower panel, ruby).

In some cases the spheres of the headplate did not seat within the kinematic features. In this case, position could not be assessed using 2P microscopy as the displacement was greater than the field of view. These events were defined as clamp failures and have been described in previous studies of voluntary head restraint (Scott *et al*., 2013). Images taken when the headplate failed to clamp could not be registered, either because the pattern of beads was completely different or there were no beads at all. For each headplate, we describe the clamp failure rate (Table 1). The average clamp failure rate for each of the headplates are as follows: ruby bearings was 3.6%, stainless steel bearings was 0.3%, machined half spheres was 12.7%. The ruby bearings and stainless steel bearings had significantly less clamp failure than the machined half spheres (p<0.001, Fisher’s Exact Test).

### Effects of implantation and stability during imaging

Next, we evaluated clamp stability by 2P imaging of populations of GCaMP6f neurons in the hippocampal CA1 region in head-fixed, transgenic mice locomoting on a wheel (Figure 5). Measurement of brain position as the animal locomoted revealed a RMS position for the stainless steel headplate of 1.13μm in the anterior-posterior (AP) axis, 0.62μm in the medio-lateral (ML) axis, and 0.16μm in the Z-axis (Figure 5C). Consistent with previous results, we found that the greatest magnitude in brain motion was observed in the AP axis during running (Figure 5E, Dombeck *et al*., 2007). The magnitude of brain motion we measured was consistent with previous results with previous headplate designs and small enough to be corrected using conventional subpixel motion correction software (Pnevmatikakis, and Giovannucci, 2017).

**Figure 5.**
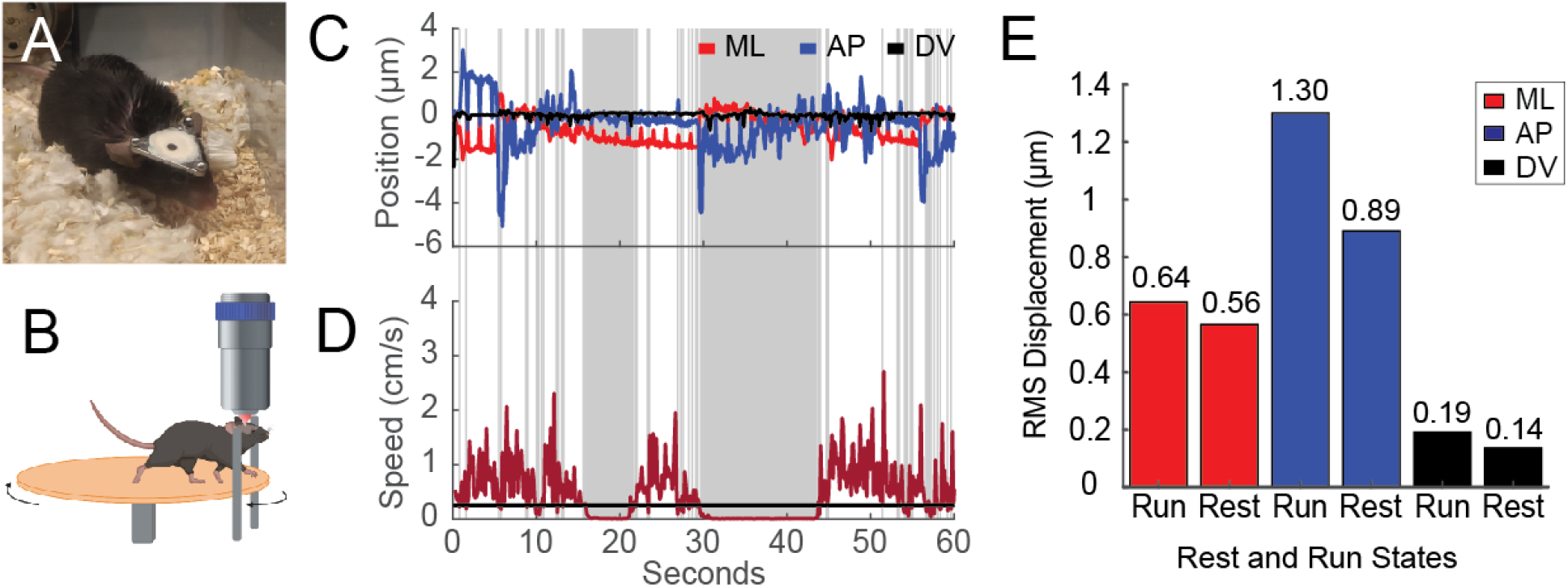
Brain stability achieved during awake behaving mouse imaging using the clamp. A) Photograph of a mouse that has been surgically implanted with the kinematic headplate. B) Diagram of the mouse on the wheel relative to the clamp and microscope. C) Evaluation of brain motion in head fixed mice on a behavioral running wheel by 2P imaging and subpixel motion correction. Blue line shows brain position as a function of time along the anterior-posterior (AP) axis, red line shows position along the medio-lateral (ML) axis, and black line shows position along the dorsal ventral axis (DV). Gray shared regions indicate periods of rest (<0.25cm/s). E) Comparison of RMS brain displacement during running (Run) and stationary (Rest) periods. The greatest magnitude in brain motion was observed in the AP axis during running, consistent with previous results (Dombeck *et al*., 2007).

Next, we sought to evaluate the presence of signal changes caused by brain motion along the z-axis. *In vivo* calcium imaging is particularly sensitive to artifacts caused by brain motion parallel to the imaging axis. As neurons move into and out of the focal plane they can cause changes in the observed fluorescence signal, which adds noise, and potentially “false positive” transients to measurements of calcium dynamics (Dombeck *et al*., 2007). To evaluate the potential presence of z-motion artifacts we imaged the same GCaMP6f+ neuronal population at 920nm and 820nm, the isosbestic point for the GCaMP family of calcium indicators (Figure 6). Fluorescence efficiency at the isosbestic point is calcium independent (Barnett *et al*., 2017) and allows evaluation of fluorescence transients caused by brain motion and other factors. We identified 163 cells at both wavelengths and compared the fluorescence dynamics between these wavelengths at the single cell-level and the population-level (Figure 6, middle and right columns). On average, neurons recorded at 920nm exhibited a significant fluorescence transient rate of 0.05Hz, (662 transients recorded total) while recording the same cells at 820nm yielded an average transient rate of 0 Hz (0 total transients recorded total), consistent with a false positive rate of <1 detected transients per 163 cells over 1.5 minutes. Our results suggest that our headplate is stable enough for multiphoton calcium imaging *in vivo* without a detectable rate of false positive (i.e. calcium independent) transients.

**Figure 6.**
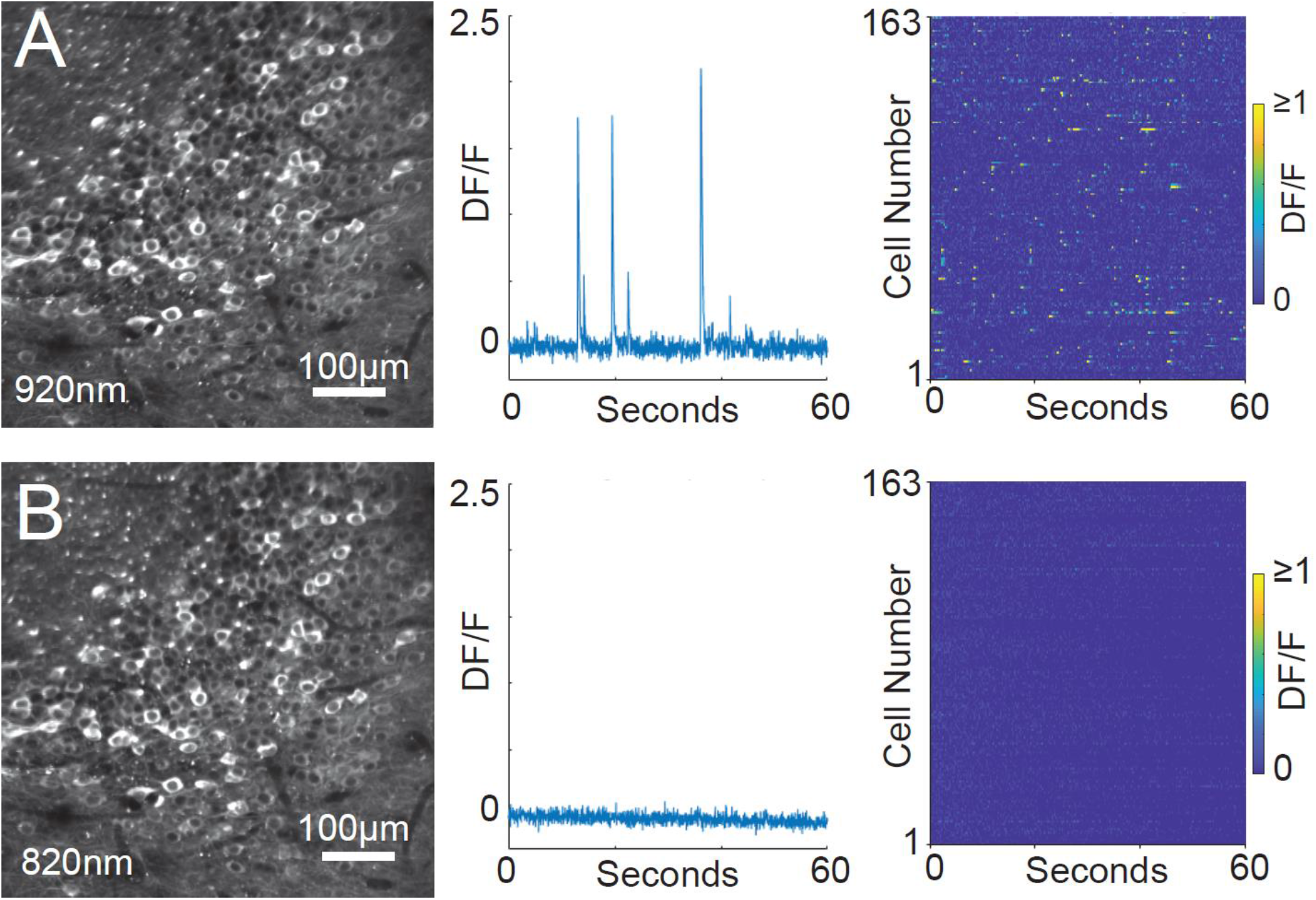
Clamp provides the stability necessary for multiphoton cellular resolution calcium imaging in awake behaving mice. A) Image of the CA1 region from a transgenic GCaMP6f animal, imaged using a 2P microscope at 920nm (left). Fluorescence trace for a single cell imaged at 920nm (middle). Population activity for the field of view in panel A at 920nm (right). Color indicated the normalized fluorescence (DF/F). B) Image of the same field of view at 820nm (left), the approximate 2P isosbestic point for the GCaMP family of calcium indicators (Barnett *et al*., 2017). Fluorescence efficiency at the isosbestic point is not calcium dependent and thus allows evaluation of fluorescence transients caused by un-corrected brain motion. Fluorescence trace for the same single cell from panel A, imaged at 820nm (middle). Population activity for the field of view in panel A at 920nm (right).

## Discussion

Here we report the development and characterization of a new head-restraint system capable of high registration accuracy, repeatability, and stability, and is small enough to be worn by mice. This system, which incorporated kinematic design features, achieved micron-scale registration repeatability and allowed stable multiphoton cellular resolution calcium imaging in mice running on a wheel with minimal motion artifacts. Furthermore, the addition of ultrahard polished ruby balls as kinematic features significantly improve the registration accuracy of the system. These results extend kinematic registration systems to mice and improve the registration accuracy of the clamp to the submicron regime.

Kinematic headplates have previously been developed for rats (Scott *et al*., 2013, Rich *et al*., 2018). These systems exhibit registration accuracy and stability sufficient for cellular resolution two-photon imaging. To achieve this performance, these systems utilized a headplate coupling-systems inspired by a Kelvin-style kinematic clamp, in which three spherical features on one side of the mount mate with a conical depression (which ideally should be a trihedron), a flat surface and vee groove. In contrast our system utilized a Maxwell coupling system, in which the spherical features mate with three vee grooves. Maxwell couplings, also known as three-groove couplings, have reduced contact stress in its couplings (if a gothic arch groove is used instead of a flat side vee groove), reduced manufacturing costs (Slocum 1995), and are known to be more repeatable than Kelvin-style couplings (Evans 1989).

Registration accuracy in kinematic coupling systems depends on several factors including the smoothness of the contact surfaces, the hardness of the materials, and the forces applied during coupling (Slocum 1992a, Slocum 1992b, Hale and Slocum 2000, Hale and Slocum 2001). We tested a range of surface contact materials (ruby bearings, 304 stainless steel bearing balls, 17-4 ph stainless steel spheres machined integral with the plate). Each contact material produced a different range of registration accuracy performance. One parameter which we did not fully explore in this study is the coupling force, which is known to influence repeatability of kinematic couplings: It should be as high as possible up to the point of inducing half of the allowable infinite life material shear yield stress in the region of highest shear below the contact surface (Hertz stress) zone (Slocum 2010).

All the material combinations tested met the stress ratio criteria, and greater clamp force could be used. The contact stress rises with the applied force to the 1/3^rd^ power (see Table 2), and holding force varied across our headplate designs. For example, ruby bearings produced less holding force than the bearings or the machined spheres because they do not help with completing the magnetic circuit, which must thus only rely on the air gap. To remedy this in the future, flux focusing rings could be provided around the magnets. We point out that coupling force is also a key determinant in the stability during in vivo imaging and in the ability of subjects to voluntarily disengage with the head fixation apparatus (Scott et al. 2013). Therefore, optimization of clamp holding force used in voluntary restraint and imaging studies should involve attention to the subject’s ability to operate the clamp and may differ depending on the species.

**Table 2.** Forces and predicted contract stress of our clamp and different contact materials.

| **Groove material** | **Tensile**<br>**yield (MPa)** | **Modulus**<br>**(GPa)** | **Poisson**<br>**ratio** | **Coupling**<br>**force (N)** | **Max shear stress /**<br>**(0.5\*Tensile yield)** | **Max force**<br>**possible (N)** |
| --- | --- | --- | --- | --- | --- | --- |
| 17-4 Ph 1025 | 1138 | 200 | 0.29 |  |  |  |
| **Ball material** |  |  |  |  |  |  |
| Ruby | 2070 | 345 | 0.21 | 0.9 | 0.338 | 2.9 |
| 17-4 Ph 1025 | 1138 | 200 | 0.29 | 2.4 | 0.407 | 4.4 |
| SS bearing balls | 1725 | 200 | 0.29 | 3.4 | 0.456 | 4.5 |

### Comparison with other systems

Various coupling systems have been developed for rodent head restraint and several have been designed to achieve reliable repositioning. Recent work from the Allen Institute describes methods for a reproducible head-fixation pipeline (Groblewski *et al*., 2020) and reports a median registration adjustment of 16μm between sessions in each the horizontal and vertical axes, small enough for researchers to manually correct to the target FOV. The Allen Institute head-fixation system reports motion while the animal is head-restrained ranging from 1-5μm, with larger movement in the z-axis towards the edge of the implant.

Other existing head-fixation systems made by laboratories (Andermann *et al*., 2010, Guo *et al*., 2014, Hao *et al*., 2021) or by companies (Neurotar, personal correspondence) achieve micrometer-scale precision or larger. These head-fixation systems implement headplates with non-kinematic (such as threaded holes and screws) or quasi-kinematic methods to achieve such precision, and thus they still suffer from micrometer-scale z-axis fluctuations, which may introduce false positive (calcium independent) transients. Using a three-groove kinematic system with sufficiently smooth and hard contact surfaces, we observed an order of magnitude improvement in z-axis repeatability compared with previous studies.

## Limitations

In typical laboratory environments, particularly those involving rodent subjects, dirt and debris can accumulate on instruments over time. Although we did not characterize this effect in our system, in principle the accumulation of debris may lead to increased clamp failure rate and decreasing registration accuracy. Regularly cleaning the vee grooves and the contact spheres on the headplate can prevent dirt from affecting registration accuracy. In addition, wear caused by repeated clamp activations could lead to changes in the smoothness of contact features. Investigators interested in maintaining high accuracy with any of the contact materials previously described can consider polishing the vee grooves in the Maxwell clamp, by using a set of square polishing stones with increasing grit and ensuring the polishing is uniform across the vee groove. Polishing can increase the accuracy by decreasing surface roughness from milling and wear (Slocum and Donmez 1988). However, such maintenance may require increased experimental oversight or decreased performance especially in chronic voluntary restraint applications.

### Experimental outlook

We anticipate that the headplate clamp system reported here can facilitate mouse experiments where high precision and stability are required, such as time-lapse developmental studies, or registration of the same subject across multiple instruments. In addition, a promising next step is incorporation of these headplates into a system for mouse voluntary head restraint. Recently several groups have reported systems to train mice in voluntary head restraint (Aoki *et al*., 2017, Murphy *et al*. 2020, Hao *et al*., 2021). In these systems mice show robust self-initiation fixations over months and can perform decision making tasks during fixation (Murphy *et al*. 2020, Hao *et al*., 2021). Such systems allow precise presentation of sensory stimuli and provide the stability needed for optogenetic stimulation and mesoscale optical imaging using widefield microscopes.

The incorporation of a kinematic clamp system, such as we describe here, would enable the stability and repeatability necessary for cellular resolution optical stimulation and imaging experiments as has been shown in rats (Scott *et al*. 2013, 2017, Rich *et al*. 2018). Such an approach may help to reduce the stress typically associated with forced head restraint (Juczewski *et al*., 2020) and could be adapted for other animal models, such as primates, which can be trained in voluntary restraint systems (Walker *et al*., 2020).

It is noted here that the semiconductor industry needed a standard interface for wafer carrying pods when it transitioned from 150mm to 200mm (and then 300mm) wafers. It settled on three-groove kinematic couplings, which use the same design spreadsheet referenced herein (Slocum 1992c, see SEMI E57 Specification for Kinematic Couplings Used to Align and Support 300 mm Wafer Carriers). The neuroscience research community may wish to consider adopting a similar three-groove kinematic interface and design method.

## Acknowledgements

We would like to thank D. Boas, A. Chen, J. Giblin, K. Kilic, and S. Park for providing technical imaging assistance. A. Monasterio performed surgical implantation of mice for the *in vivo* imaging experiment. T. Otchy provided feedback throughout the design process. A. Krupp assisted with the scanning electron microscope and provided feedback on the related sections, and J. Javor assisted with the optical profilometer.

The MIT Precision Engineering Research Group provided design feedback. D. Campbell, J. Estano, and B. Sjostrom of the BU Engineering and Product Innovation Center, and G. Thayer of the BU Scientific Instrumentation Facility provided design feedback on early prototypes. We would like to thank BioRender for the illustration of the mouse on the wheel. This work was supported by a NARSAD Young Investigator Award from the Brain and Behavior Research Foundation to BBS.

## Contributions

S.J.K. and B.B.S. conceived the presented idea. A. H. S. guided the use of the kinematic coupling design spreadsheet and provided design and manufacturing feedback. S.J.K designed and fabricated all parts and took the lead on shaping the findings. S.J.K. and B.B.S. designed the 2P imaging experiments. S.J.K. conducted all experiments, processed and analyzed data, drafted the manuscript, and designed the figures. S.J.K and B.B.S. interpreted the results and wrote the manuscript. B.B.S. supervised the overall direction and planning of the project.

## Supplemental Material

**Supplemental Figure 1.**
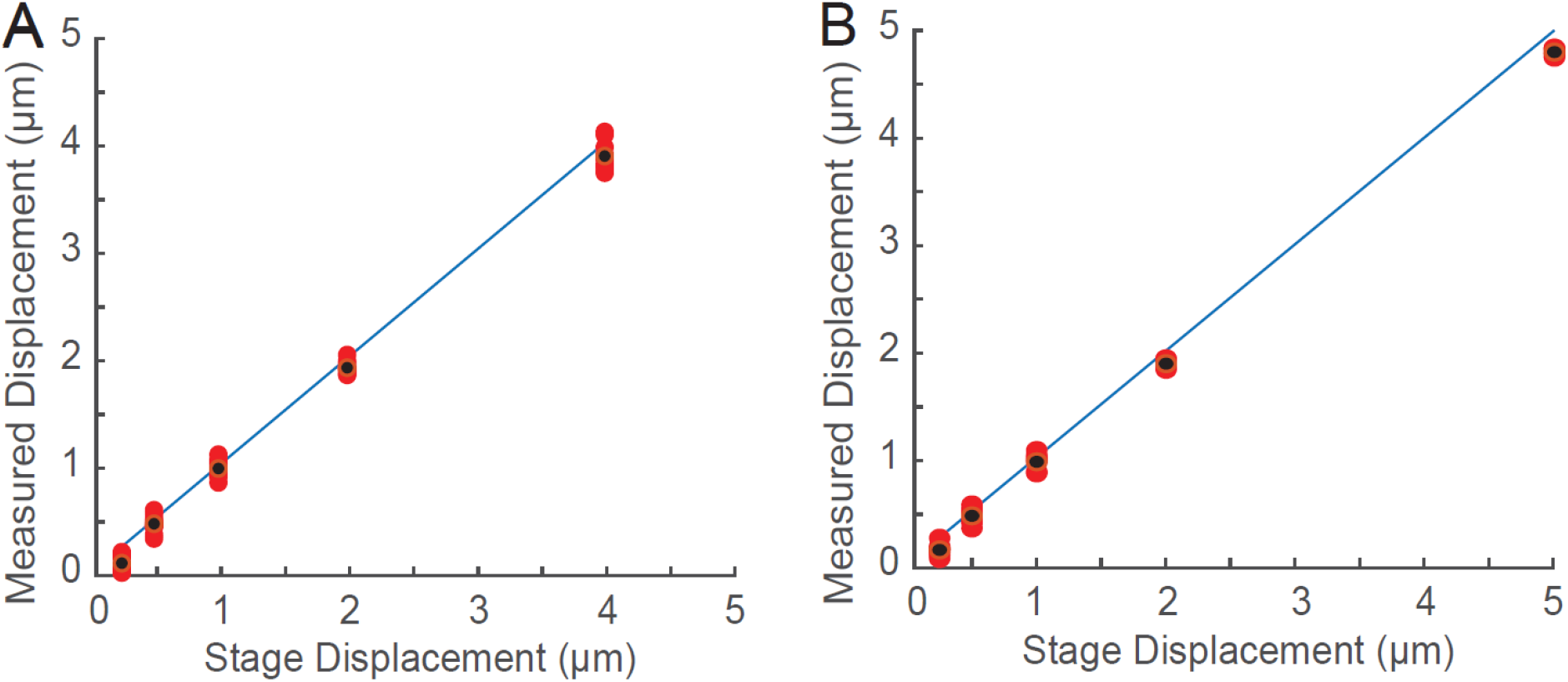
Evaluation of displacement measurement accuracy. To assess our accuracy in measuring displacement, we commanded the microscope stage to move a known distance and took an image after each movement. Each red dot is the new stage position subtracted from the previous stage position (defined as displacement, see methods), plotted against the measured displacement from image analysis. Black dots indicate the mean displacement for each condition and the blue line is the unity line (y = x). A) Image displacement after commanding the microscope objective to move in the horizontal direction: 4μm, 2μm, 1μm, 0.5μm, 0.25μm. The x-axis shows the commanded stage displacement, and the y-axis shows the displacement as measured by a motion correction algorithm. B) Image displacement after commanding the microscope objective to move in the vertical direction 5μm, 2μm, 1μm, 0.5μm, and 0.25μm.

